# Quantifying Metagenomic Strain Associations from Microbiomes with Anpan

**DOI:** 10.1101/2025.01.06.631550

**Authors:** Andrew R. Ghazi, Kelsey N. Thompson, Amrisha Bhosle, Zhendong Mei, Yan Yan, Fenglei Wang, Kai Wang, Eric A. Franzosa, Curtis Huttenhower

## Abstract

Genetic and genomic variation among microbial strains can dramatically influence their phenotypes and environmental impact, including on human health. However, inferential methods for quantifying these differences have been lacking. Strain-level metagenomic profiling data has several features that make traditional statistical methods challenging to use, including high dimensionality, extreme variation among samples, and complex phylogenetic relatedness. We present Anpan, a set of quantitative methods addressing three key challenges in microbiome strain epidemiology. First, adaptive filtering designed to interrogate microbial strain gene carriage is combined with linear models to identify strain-specific genetic elements associated with host health outcomes and other phenotypes. Second, phylogenetic generalized linear mixed models are used to characterize the association of sub-species lineages with such phenotypes. Finally, random effects models are used to identify pathways more likely to be retained or lost by outcome-associated strains. We validated our methods by simulation, showing that we achieve more accurate effect size estimation and a lower false positive rate compared to alternative methodologies. We then applied our methods to a dataset of 1,262 colorectal cancer patients, identifying functionally adaptive genes and strong phylogenetic effects associated with CRC status, sometimes complementing and sometimes extending known species-level microbiome CRC biomarkers. Anpan’s methods have been implemented as a publicly available R library to support microbial community strain and genetic epidemiology in a variety of contexts, environments, and phenotypes.

## Introduction

Genetic variation within bacterial species can lead to dramatic differences in functional potential, with corresponding impacts on host health or environmental phenotypes. For example, strains of *Escherichia coli* can be enteric pathogens (e.g. Shiga-toxigenic strain O104:H4)[1] or increase risk of chronic disease (e.g. inflammatory bowel disease, IBD, or colorectal cancer)[2, 3]. However, apparently benign strains of *E. coli* are prevalent in “healthy” gut microbiomes[4–6], and strains such as *E. coli Nissle* 1917 are generally regarded as probiotic[7, 8]. Similarly, *Ruminococcus gnavus* contains at least two sub-species lineages, only one of which carries functionality associated with IBD and rheumatoid arthritis[9, 10]. Thus, the ability to accurately identify and characterize bacterial strains in population-based studies is critical in understanding the causal links between the microbiome and human disease. This is particularly true given the rapidity with which microbial genetics change and evolve, yielding new subspecies-level variation in almost every setting.

The definition of what constitutes a “strain” varies across scientific contexts[11]. Originally strains were defined in terms of cultures originating from a single colony or cell [12], though this usage has declined over time with culture-free methods of isolating and characterizing strains using metagenomic sequencing[13, 14]. Using such methods, a “strain” can be defined as a set of fully clonal (i.e. identical) genomes, but this is complicated by the presence of a spectrum of variants ranging from minor (e.g. a single nucleotide polymorphism) to substantial (e.g. horizontal gene transfer within and across species). It can thus be useful to speak in terms of “subspecies population structure,” clades, or lineages instead of “strains” when species-wide phylogenetic relatedness is more directly relevant to the experiment at hand. Here, we will operationally define strains as “a set of cells or genomes within a narrow, monophyletic subspecies clade,” which in practice will fall within a greater than 99% genomic nucleotide identity threshold (in contrast to typical species-level definitions within a 95-97% identity range) [15, 16].

Several methods have emerged for strain profiling from metagenomes based on similar definitions[11], including StrainGE[14], StrainPanDA[17], StrainPhlAn[18], and PanPhlAn[19]. Such methods typically operate on one of two principles: comparing reads to reference or assembled genomes for variant calling (often single nucleotides)[11] or comparing gene profiles to identify gains and losses[20] (“structural variants”). They must overcome challenges such as ambiguous read assignment across closely-related taxa and highly variable genome coverage, but as evidenced by the diversity of methods now available, this has become increasingly practical. However, downstream inferential methods for quantifying the epidemiological connection between strain characteristics and other variables (such as host health or environmental outcomes) are comparatively underdeveloped.

This methodological gap between the ability to identify strains and the ability to understand what those strains are doing emerges from several problematic characteristics of strain data. Strain data are noisy, high-dimensional, frequently zero-inflated, and commonly exhibit complex phylogenetic relatedness. These problems emerge partly from the complexity of microbial community genetics themselves, and partly from the ever-increasing catalog of features that can be quantified from metagenomic sequencing. If one defines a strain as “a unique set of microbial genes”, dimensionality issues quickly ensue. For instance, metagenomic studies can quantify thousands of microbial genes[21] for each of hundreds of species in thousands of human-derived samples; these are in turn drawn from individuals who may vary in disease status, geographic location, demographics, or other relevant covariates. Detection and quantification of such genes is also made noisy due to variation in genomic coverage, both biological (e.g. origins of replication) and technical (e.g. GC bias). Finally, strains are frequently sample-specific. The more restrictive a definition of “strain” that is employed, the more likely it is to be found in only a single individual or environment, which in turn means that estimating the effects of the strain across repeated, independent samples is impossible. For this reason, strain-level statistical modeling requires methods that can aggregate subspecies structure across samples in some way, either genomically (i.e. by inspecting unique genetic elements that recur across individuals) or phylogenetically.

This combination of biology and measurement strategies suggests three complementary models by which microbial community strain epidemiology can be investigated. A phenotype may be influenced by the presence of individual genes, the abundance of aggregated gene pathways, or the net effect of all phylogenetic variation across the entire breadth of the species. Here, we present Anpan, a collection of statistical methods aimed at quantifying such effects of strain variation on outcomes in the context of microbial communities (**Fig. 1**). The corresponding three components of Anpan are 1) a gene modeling framework that uses quality control and adaptive filtering steps to identify individual microbial genes whose presence is associated with binary or continuous outcome variables (such as disease state or BMI); 2) a phylogenetic generalized linear mixed model framework that can be used to identify cases where within-species phylogeny covaries with outcome; and 3) a mixed effects model of gene pathways, which identifies pathways for which frequency of within-species carriage differs substantially among experimental groups. We validate the models by simulation, assessing the accuracy and precision of estimates of simulated effects. We then apply the models to a multi-cohort study of colorectal cancer (CRC) and identify a general pattern of increased transposase prevalence in CRC cases, as well as five species where phylogenetic structure associates with CRC risk. Anpan is available as an open-source R package that automatically handles filtering, model fitting, visualization, and summarization for each model type.

**Figure 1:**
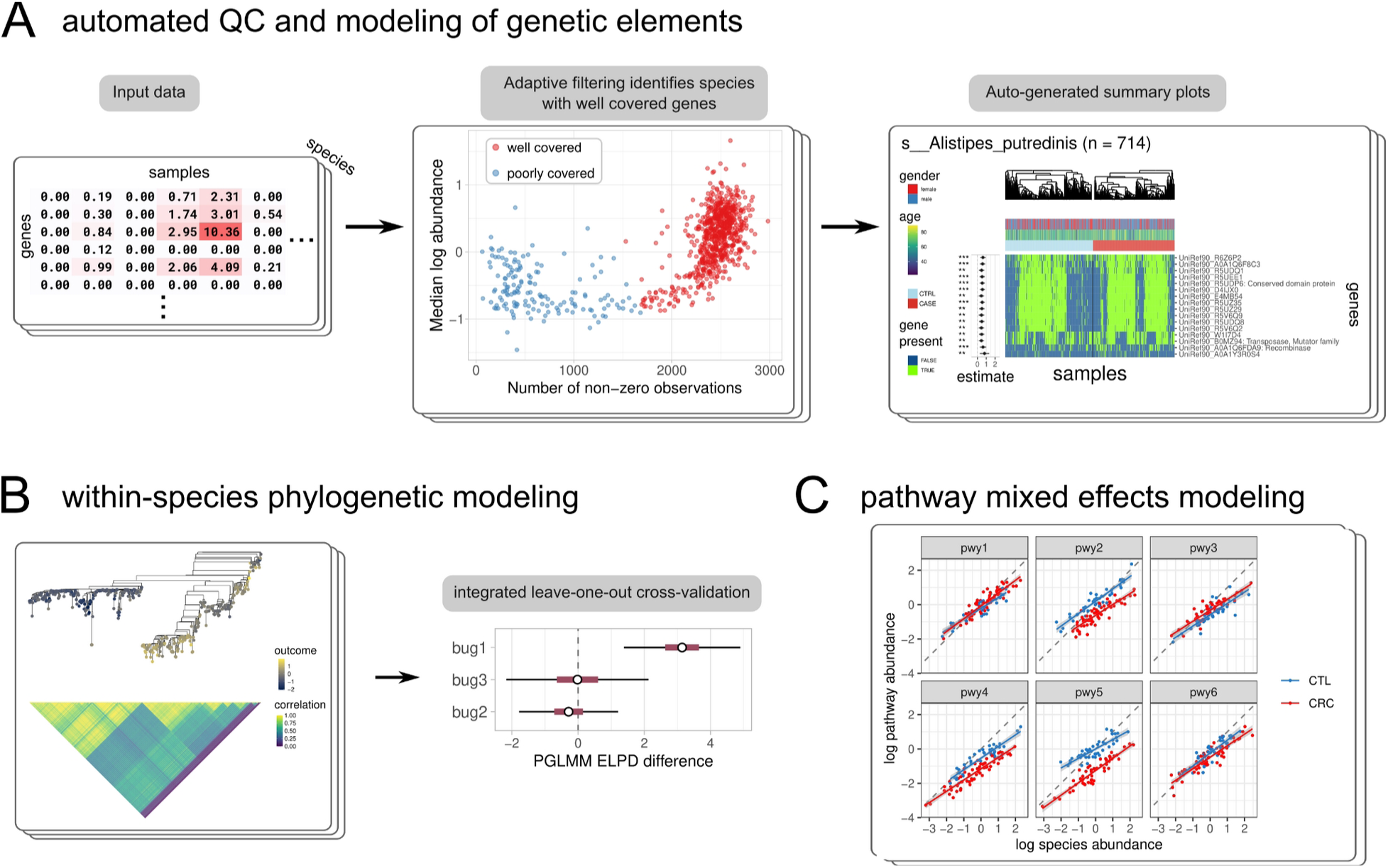
Anpan provides statistical models for strain gene carriage, phylogeny, and pathway carriage. **A)** One way in which strain variation in microbial communities can be associated with phenotypes is via the presence or absence of individual genes. Anpan’s gene-level model tests this quantitatively after an adaptive filtering step that identifies samples with sufficient data (i.e. abundance and sequencing depth) to test. **B)** Phylogenetic generalized linear mixed models are used to identify species where overall strain phylogeny across samples strongly associates with an outcome, which is quantified by leave-one-out predictive performance. **C)** Anpan’s pathway model identifies pathways (such as pathways 2, 4, and 5 in this simulated example) whose frequency of carriage within an individual species differs substantially between subject groups while accounting for species abundance. That is, pathways which are preferentially retained or lost with respect to a phenotype by strains of each species.

## Results

We validated the efficacy of Anpan’s statistical models for strain epidemiology using both synthetic data and applications to over 1,200 CRC metagenomes, the latter newly-identifying mobile elements unique to lineages of *Streptococcus parasanguinis* and *Clostridium bolteae*. First, we used simulations to demonstrate the effectiveness of our three strain analysis models, showing that our methods can reliably detect metagenomic statistical associations emerging from subspecies-level differences. We show that our method for adaptive filtering of input metagenomic profiles distinguishes samples where genetic elements from a given species are sufficiently well or poorly covered for reliable analysis. The outputs of the filtering step feed into the functional association model, which identifies associations between microbial genes and outcome variables of interest. Next, phylogenetic generalized linear mixed models (PGLMMs) are used to identify and visualize patterns of relatedness within species that explain outcome variables. Finally, pathway-level random effects models are used to identify pathways within species that vary in abundance between experimental groups.

### Adaptive filtering based on gene profiles identifies species-sample pairs with sufficient coverage for strain analysis

Very shallow shotgun metagenomes can be sufficient for species- or genus-level taxonomic profiling[22], but substantially greater depth is needed for functional or genetic (i.e. strain) profiling. For strains of a given species to be assessed within a particular metagenome, species abundance and sequencing depth must combine to provide sufficient coverage depth for gene or single nucleotide variant detection[19, 23]. Even with appropriate coverage, reliable variant calling can be challenging due to regional conservation and homology among species, or the presence of multiple strains within a species. These challenges necessitate a filtering strategy that can identify which samples can provide information on the effects of which species and which subsets of their pangenomes.

In particular, adaptive filtering of samples is necessary to identify those in which a species is absent or insufficiently covered by the available sequence depth, so that these species-sample combinations may be removed before analysis. To this end, we utilize the overall gene profile of each species in each sample to classify the species as “well covered” or “poorly covered” in that sample. Anapn first computes two statistics for each species-sample combination: the number of non-zero observations and the median log abundance of the nonzero measurements. This is analogous to a collector’s curve, in which increasing coverage eventually saturates the number of genes carried by a typical strain of that species. When visualized as a scatterplot with a point for each sample, samples with well-covered species present clearly separate from samples where the species was not present. Applying the k-means algorithm to this distribution of the two aforementioned statistics gives a useful binary labeling of “well covered” or “poorly covered” (**Fig. 2A**). Anpan retains only these well-covered species-sample pairs for the downstream modeling step. An additional diagnostic visualization (**Fig. 2B**) that plots the ranked log abundances of genes for each sample helps visually confirm the separation of well and poorly covered samples. This strategy is also highly analogous to that used by PanPhlAn to detect well-covered pangenomes[19].

**Figure 2:**
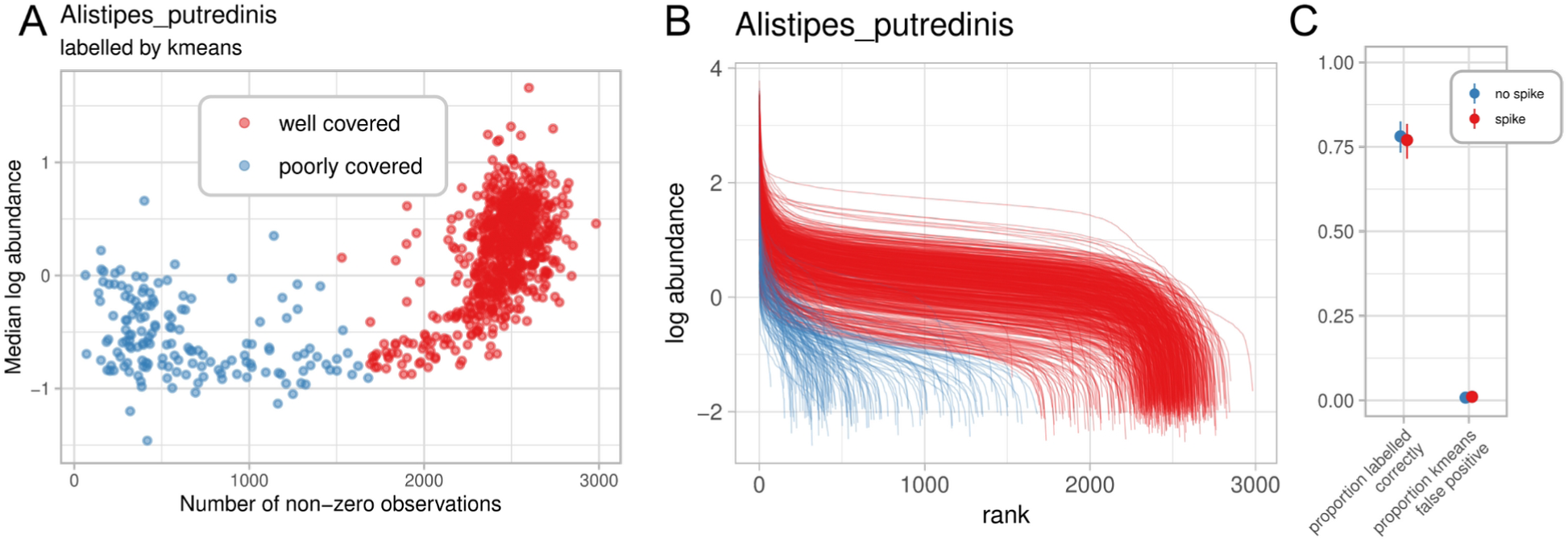
Adaptively filtering species-metagenome pairs with insufficient depth of coverage for strain profiling. **A)** Anpan, like PanPhlAn[19], relies on saturation of a species’ expected genome-wide gene count per sample as a measure of sufficient coverage. Data on gene count (i.e. breadth of coverage) by abundance (i.e. depth of coverage) is k-means clustered with k=2, grouping saturated vs. unsaturated metagenomes per species. Here, results on UniRef90 functional profiles from HUMAnN [25] in *Alistipes putredinis* are shown from the subset of 869 CRC patients in which the species was detected (of 1,262 total, **Methods**). **B)** Rank-ordered per-sample gene abundances (as per PanPhlAn) labeled as clustered by k-means. Each line is a sample, and the height of each line represents the log abundance of the i-th UniRef90 in that sample. Samples with high abundance of many *A. putredinis* genes are correctly detected as well-covered, while those with erroneously high abundance of a few genes or low abundance of an intermediate number of genes are poorly covered. **C)** Under repeated simulation, the proportion of points labelled correctly quantifies the performance of the k-means filter, showing that usually the large majority of points are labelled correctly. The proportion of false positives from the kmeans filter is generally close to 0, demonstrating that the filter generally errs on the side of being conservative, as desired.

To assess the efficacy of the resulting method, we simulated metagenomic functional profiles that correlated with risk of a binary disease status. Gene abundances were modeled as a function of simulated species abundance (**Methods**), which were in turn simulated using SparseDOSSA 2[24]. Simulated species with 1,000 genes had either 0 or 10 risk genes that increased the log odds of a simulated disease state by 1, and the abundance of the species overall increased log odds of the disease state by 0 or 1. For each condition, we generated 100 simulated experiments each with 200 controls and 200 cases. As a challenge to the adaptive filtering step, 25% of subjects were randomly selected as having “poorly-covered” genes for the simulated species, which reduced the average log gene abundances by 4. The adaptive labeling generally performed well, averaging around 80% of points correctly labeled (**Fig. 2C**, **S1**).

**Figure S1:**
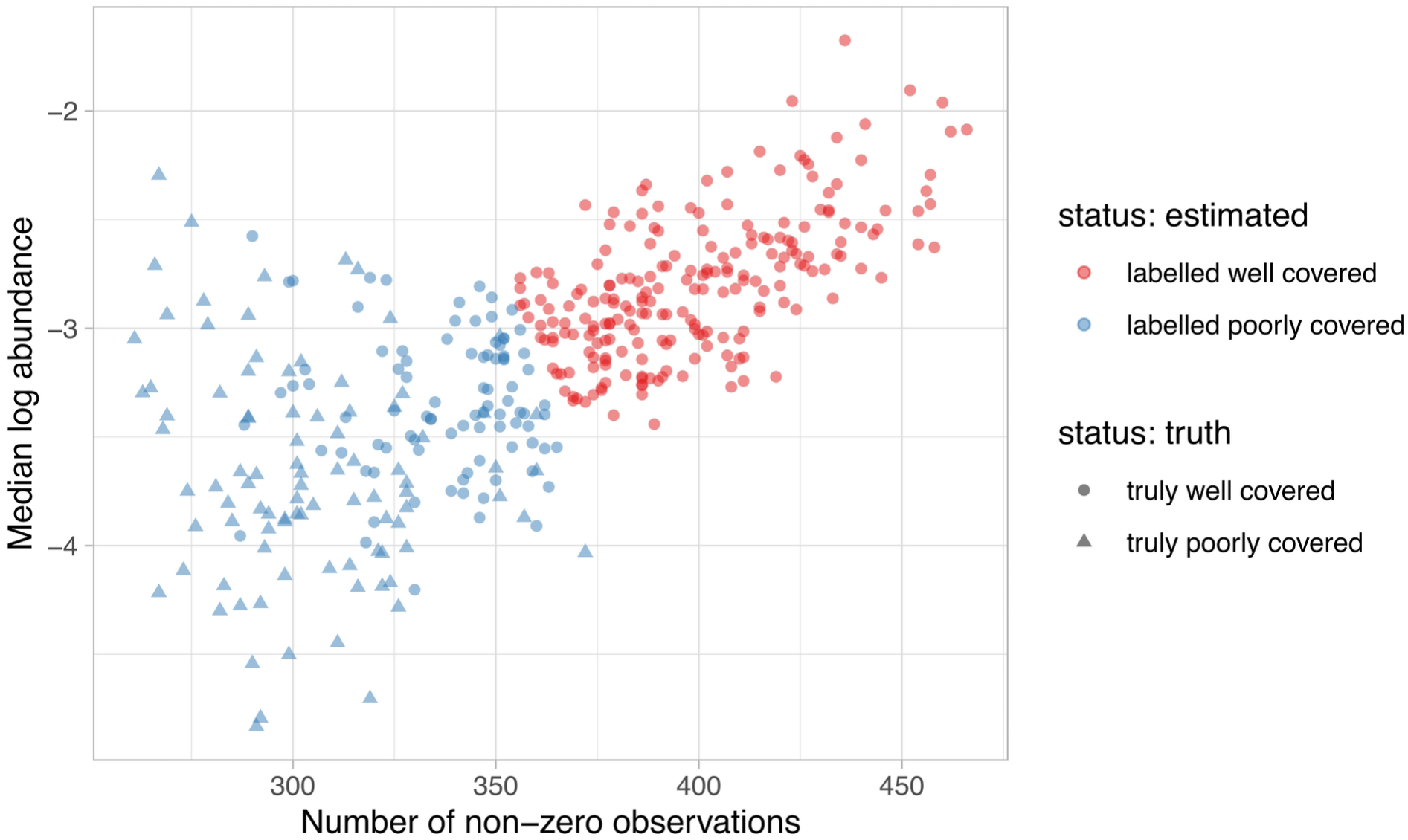
Adaptive filtering of simulated data shows accurate, conservative labeling of samples. The k-means filter applied to simulated data, alongside the true simulated status of each sample. While there are false negatives included in this example, there are no false positives as well, indicating that for these simulation parameters, the filter errs on the side of being conservative.

### Anpan accurately associates strain gene carriage with phenotypes and covariates

When it is sufficiently well-powered, associating the presence or absence of strain gene carriage with a phenotype can provide strong clues toward mechanistic interpretation[9, 10, 19, 20]. To assess the relationship between outcome variables and the presence of these genes, we use linear models with one of two methods for accounting for the multiplicity of the microbial genes. By default, Anpan fits a standard GLM of the form `outcome ~ <COVARIATES> + gene_presence` (with an appropriate link function) for each gene, where `<COVARIATES>` represents terms for additional user-specified covariates (e.g. age, sex, temperature, pH, etc.) The p-values for the `gene_presence` terms are then adjusted using the Benjamini-Hochberg method for multiple tests[26] and called as “hits” by a Q < 0.05 threshold.

This model is simple and effective, but since it tests each gene independently, it can sacrifice power as a result. Alternatively, a linear model that accounts for all genes simultaneously i.e. `outcome ~ <COVARIATES> + gene1_presence + gene2_presence + …` may be fit with the `model_type = “horseshoe”` option. This places a regularized horseshoe prior[27] on the coefficients for individual genes with (by default) an *a priori* expectation of 1% non-zero coefficients. This provides more straightforward model interpretation and can help mitigate correlated gene presence patterns, at the expense of increased computational requirements. With the horseshoe model, hits are identified by a configurable effect size threshold and 99% posterior interval excluding 0. Either model results in a list of zero or more genes per species that are differentially carried (i.e. present or absent within strains) with respect to the desired outcome phenotype and optional covariates (**Fig. 1A**). Alongside the central heatmap are intervals displaying the size and significance of the coefficients, along with color bars displaying any additional covariates.

In order to validate the performance of the gene carriage model, we evaluated it on the filtered results from the simulated metagenomic datasets produced from the previous section. We introduced an artificial effect of gene carriage for five out of 1,000 simulated genes. This effect increased the log odds of a binary outcome variable by one. In order to understand the model’s behavior in scenarios where the outcome is correlated with overall species abundance on top of any effects from individual genes, we tested this simulated effect in two scenarios: one where there was a spiked association between overall species abundance and the outcome and one with no spiked association. Together, these simulation parameters create experimentally realistic datasets where true gene effects are comparatively modest and rare and hence correspondingly difficult to detect, particularly with a binary outcome variable. Nonetheless, the model accurately utilized the sparse information provided to it, as judged by several classification metrics including an (**Fig. 3**).

**Figure 3:**
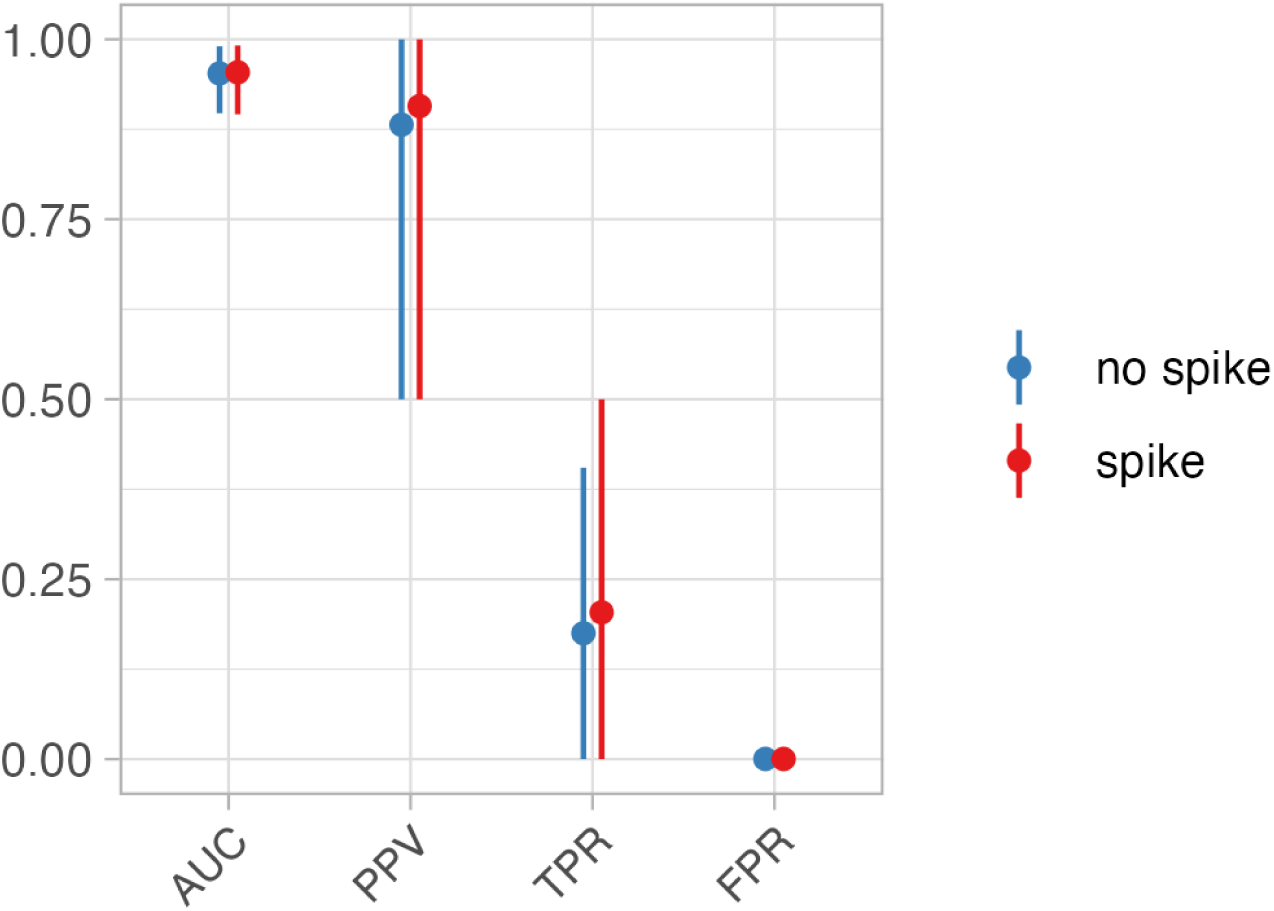
Gene carriage variants are accurately associated with simulated phenotypes. We evaluated the efficacy of the gene carriage model using the same simulated, filtered datasets derived from SparseDOSSA 2 [24], with and without spiked associations with overall species abundance that increased the log-odds of an artificial binary covariate. The Anpan gene model achieved high area under ROC (AUC) and positive predictive values (PPV) while, crucially, maintaining near-zero false positive rates. Corresponding true positive rates are modest due to the inherently weak signal and extremely high-dimensional context, i.e. thousands of genes in hundreds of species with variable (simulated) sequencing depth and effect sizes. Points and bars represent simulation means and 90% intervals, respectively.

### Detecting phenotype-associated phylogenetic lineages

Next, to address the challenge of connecting phylogenetic information to outcomes, we used phylogenetic generalized linear mixed models (PGLMMs) to model outcome variables as a function of phylogeny (**Fig. 1B**). Briefly, PGLMMs are a subclass of random effects models where each observation is allocated its own random intercept. The correlation between any pair of individual random effect terms is related to the branch length between them in an input phylogenetic tree. In Anpan’s setting, one tree is constructed per species, in which each tip represents the dominant strain of that species within one metagenomic sample. Leaves (i.e. equivalently, strains or samples) that are adjacent in the tree thus induce highly correlated random effect terms, while leaves that are further apart have lower correlation. With an identity link function, this model can be represented as:

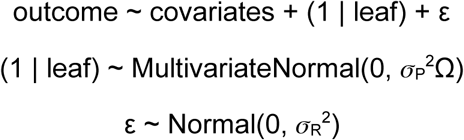

Where (1 | leaf) represents the vector of “phylogenetic effects” on each leaf, σ_P_ is the “phylogenetic noise”, Ω is the correlation matrix derived from the tree, ε are the residuals, and σ_R_^2^ is the residual variance.

While this results in a high-dimensional model that uses a parameter for every observation (in addition to the usual regression coefficients of other covariates and noise terms), the tree-derived correlation matrix imposes a correspondingly high number of constraints on the posterior distribution. In order to identify species with nonrandom phylogenetic assortment with respect to a phenotype, Anpan uses Stan[28] to estimate the parameter values and leave-one-out expected log pointwise predictive density (ELPD)[29]. The latter provides a model comparison criterion to compare the PGLMM model’s fit relative to a “base” model without the phylogeny term. The ELPD is conceptually similar to the common Akaike information criterion (AIC [30]), but provides an intuitive leave-one-out interpretation, standard errors, and additional computational diagnostics that help detect potentially inaccurate estimates. The flexibility of the phylogeny term can cause instability in the point-wise log-likelihood values, therefore we further implemented the integrated importance sampling approach described in[31] to obtain importance weights when computing the ELPD for Anpan.

In order to quantify the efficacy of the PGLMMs for the purpose of within-species phylogenetic lineage association, we performed a simulation study (**Fig. 4**) that varied sample size, phylogenetic noise (σ_P_), and residual noise (σ_R_). We compared the parameter estimates and classification statistics from PERMANOVA[32] (as implemented in vegan::adonis())[33], MiRKAT[34], and two types of PGLMMs available in Anpan, unregularized and regularized. Because the data are simulated with the causal flow from phylogeny to outcome (and not the opposite direction), the permutational linear methods dramatically underestimate the R^2^ statistic. This speaks to the under-appreciated fact that under a structure where phylogeny acts as a cause of the outcome, “proportion of variance explained” is an incorrect interpretation of the PERMANOVA R^2^ statistic. In comparison, unregularized PGLMMs are generally anti-conservative at low sample sizes, overestimating R^2^ due to their flexibility with the default weak priors.

**Figure 4:**
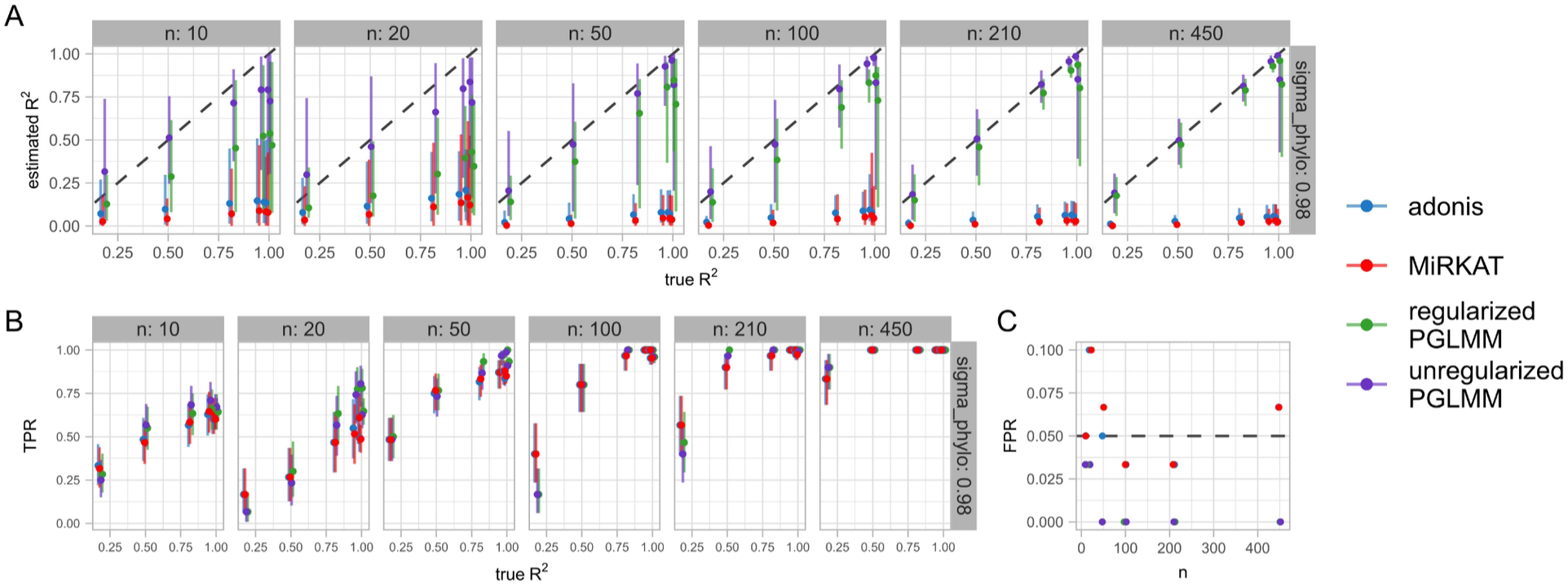
Regularized PGLMMs identify within-species phylogenetic signal impacting phenotype. Across a variety of simulated datasets with varying sample size, phylogenetic noise (σ_P_), and residual noise (σ_R_), phylogenetic models outperform distance-based permutational linear models in identifying species with phenotype-associated lineages. In all plots, points have slight horizontal shift to prevent overlap. **A)** Estimated R^2^ values for phylogenetic effect (mean and 95% interval across simulations) against true R^2^ values driving the simulation at σ_P_ = 0.98 (see **Fig. S2** for other values of σ_P_). Notably, permutational linear models consistently and dramatically underestimate R^2^. Unregularized PGLMMs overestimate R^2^ at very low sample sizes, which is alleviated by the regularized PGLMM. **B)** True positive rates across simulations. PGLMMs generally exhibit higher power at lower n, though permutational linear models eventually catch up with increasing sample size. **C)** False positive rates across simulations. PGLMMs with default settings show an elevated FPR at the lowest sample sizes before dropping to zero by n = 100. Permutational linear models consistently yield roughly 5% FPR, consistent with their α = 0.05 significance threshold.

Because we cannot know the scale of the phylogenetic noise nor the residual noise *a priori*, Anpan thus defaults to a regularized PGLMM that imposes a penalty term on the ratio of the noise parameters: (σ_P_ / σ_R_) ~ Gamma(1,2). One interpretation of this regularizing prior term is that *a priori* there is an 86% chance that this ratio is less than 1, with values closer to 0 weighted more highly. In practice, the regularization dramatically reduces the variance at low sample sizes, while only imposing a small negative bias at larger sample sizes (**Fig. 4B-C**). Notably the PGLMMs show generally higher true positive rates (with the regularized model about 1% higher than the unregularized), though the alternatives eventually catch up with increasing sample size. The false positive rate of PGLMMs is inflated at very low sample sizes (n < 20, though this effect could be mitigated by altering the weak-by-default priors), but drops to zero beyond that, while the permutational linear models continue to hover around their nominal α level of 0.05.

**Figure S2:**
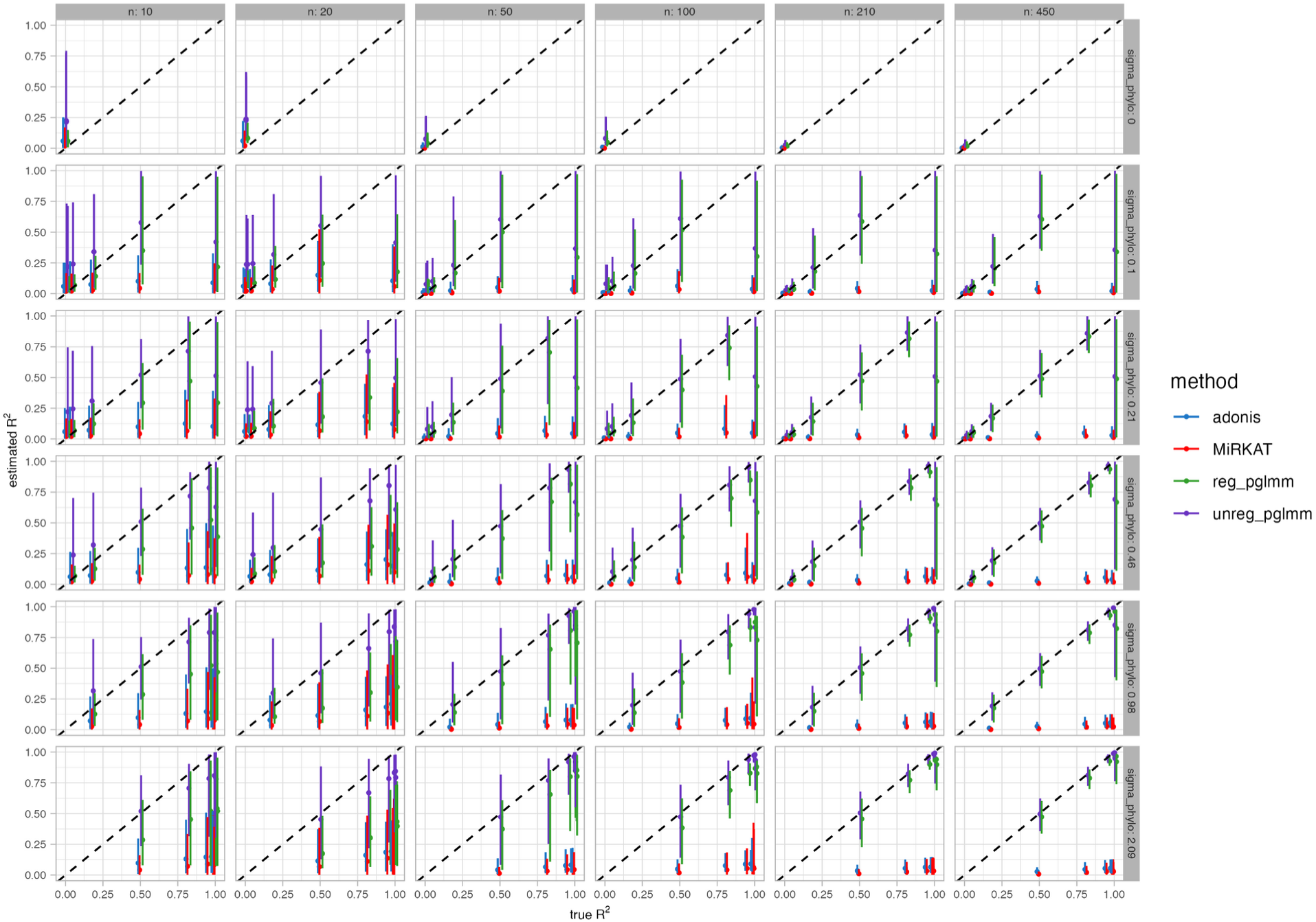
Simulated R^2^ estimates with varying σ_P_. Extended results from Fig. 4A. Permutational linear models consistently underestimate R^2^ across all values of σ_P_.

### Random effects models identify pathways preferentially retained or lost among strains with respect to phenotype

Carriage of microbial pathways presents an alternative approach to quantifying strain level effects in metagenomic data. While less genetically-specific than functional profiles based on individual gene families (e.g. UniRef90s), pathways that represent jointly functional (and thus gained or lost) genes (e.g. from MetaCyc[35]) can provide an interpretable view of microbial metabolism. Given that species-stratified pathway levels (again provided by profiling tools like HUMAnN [36]) are strongly correlated with overall species abundance, pathways carried at lower levels by disease-associated strains accumulate at a different rate with increasing species abundance. When visualized on a log-log scale of species versus pathway abundance (**Fig. 1C**), this behavior is captured as a vertical shift between phenotype groups.

This biological behavior lends itself to a random effects model, which provides several benefits in identifying pathways preferentially retained or lost among strains. First, the relationship between species abundance and pathway abundance is estimated globally, using data from all pathways in the species. This allows disaggregation of the species’ overall pathway carriage from the frequency with which it is absent in individual strains. Second, pathway-specific random intercepts allow variation in the baseline pathway abundance (i.e. within-species copy number), but the distribution of intercept estimates are shrunken across pathways. Finally, another pathway-specific random group effect term allows for estimates in between-phenotype variability in pathway carriage. Pathway-specific random effects (for both the intercepts and group effects) naturally provide adaptive regularization, while also allowing individual pathway terms to be estimated accurately when the data provide sufficient evidence. An Exponential(3) prior on the standard deviation of group effects provides relatively aggressive shrinkage on the parameters of interest, while the remaining priors on the other auxiliary parameters are deliberately weak in comparison (**Methods**). Ultimately, each pathways are called as binarized “hits” by three criteria: the 98% posterior interval of the pathway’s group effect excludes 0, the absolute value of the pathway’s group effect exceeds 0.25 (i.e. a 1.78-fold increase or decrease), and the estimate of the overall pathway-species linear coefficient must be positive.

To assess the performance of this model on biologically realistic data, we simulated pathway abundances as a function of species abundances again drawn from SparseDOSSA 2[24]. From the species abundances, we randomly simulated pathway abundances by running the generative structure of the model in reverse. For each simulation, we constructed a random profile per species with thirty pathways, of which three had a true, nonzero difference in abundance between binary phenotype groups. This difference was varied between zero and one, while the noise about the linear relationship was maintained at one. This provides simulated, biologically-realistic datasets of pathway profiles with varying degrees of signal. By applying the pathway model to these datasets, Anpan was able to accurately detect and classify associations above a minimal detectable level under real-world conditions (**Fig. 5**).

**Figure 5:**
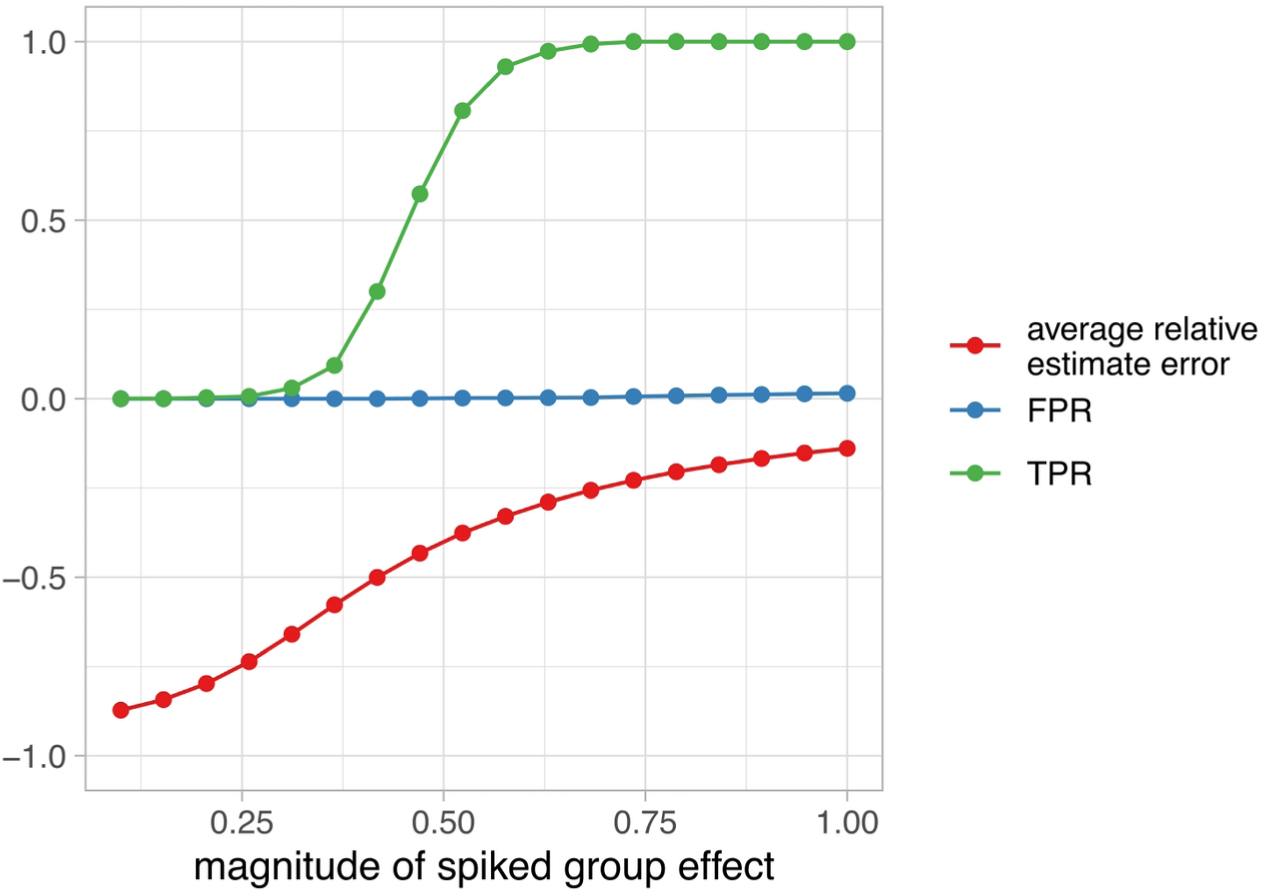
A random effects model captures phenotype-linked differences in pathway retention among simulated strains. We simulated pathway profiles correlated with species abundance using SparseDOSSA 2[24] and the generative structure of the model (**Methods**) These simulations included a species-level association and varying phenotype effect across simulated pathways. After fitting the model, we assessed the results in terms of relative error, true positive rate, and false positive rate. The x-axis represents the log10 fold difference in pathway abundance between the two phenotypic groups. Across a range of effect sizes, realistically-simulated pathway datasets showed TPR gradually increasing as average relative error of the group effect estimates gradually decreased. At high spiked effects, rare false positives became slightly more frequent due to the difference in model specification between the inference and simulation frameworks.

### Anpan identifies genetic mobility and subspeciation in CRC

As an initial driving biological problem, we applied each of Anpan’s three main models to a pooled set of eight CRC studies totaling 1,262 patients (662 controls and 600 CRC cases)[37–44]. Using UniRef90 profiles from HUMAnN 3, the gene model identified 5,198 gene families across 85 species significantly differential with respect to CRC status (i.e. Q < 0.1). Thirteen of these species (*Akkermansia muciniphila, Alistipes finegoldii, Alistipes putredinis, Bacteroides caccae, Clostridium symbiosum, Dorea longicatena, Eubacterium sp CAG 180, Faecalibacterium prausnitzii, Holdemanella biformis, Oscillibacter sp CAG 241, Paraprevotella xylaniphila, Ruminococcus torques,* and *Tyzzerella nexilis*) exhibited suggestive evidence of confounding phylogenetic structure, yielding Q < 0.1 for over 1% of their genes tested. Recent studies have identified multiple species level lineages in several of these clades[45–47], supporting that broad, functionally distinct phylogroups within a species can introduce many collinear genes with statistically indistinguishable effects. Setting aside these species, we were left with 1,460 genes in 72 species associated with CRC status with FDR Q < 0.1.

The strongest result, UniRef90_A0A379CDB2 in *Streptococcus parasanguinis* (coefficient = 1.71 +/− 0.29, Q = 4.6 × 10^−4^, **Fig. 6A**), is a transposase, as are 19 of the other top results across 13 other species. To some degree, these genes serve “true positives,” as we would expect differences in microbial strains to be particularly enriched for mobile genetic elements. One possible explanation of the enrichment of this function among our significant gene-CRC associations is that these genes help microbes acquire other genetic elements necessary to adapt to the unusual microenvironment of the CRC gut, possibly indicating a common survival strategy. A similar type of result is UniRef90_A0A1E3A1S6 in *Clostridium bolteae* (coefficient = 1.33 +/− 0.29, Q = 2.8 × 10^−3^, **Fig. S3**), a DNA topoisomerase, which under this hypothesis could be performing a related role by modifying DNA topology to allow integration of mobile elements.

**Figure 6:**
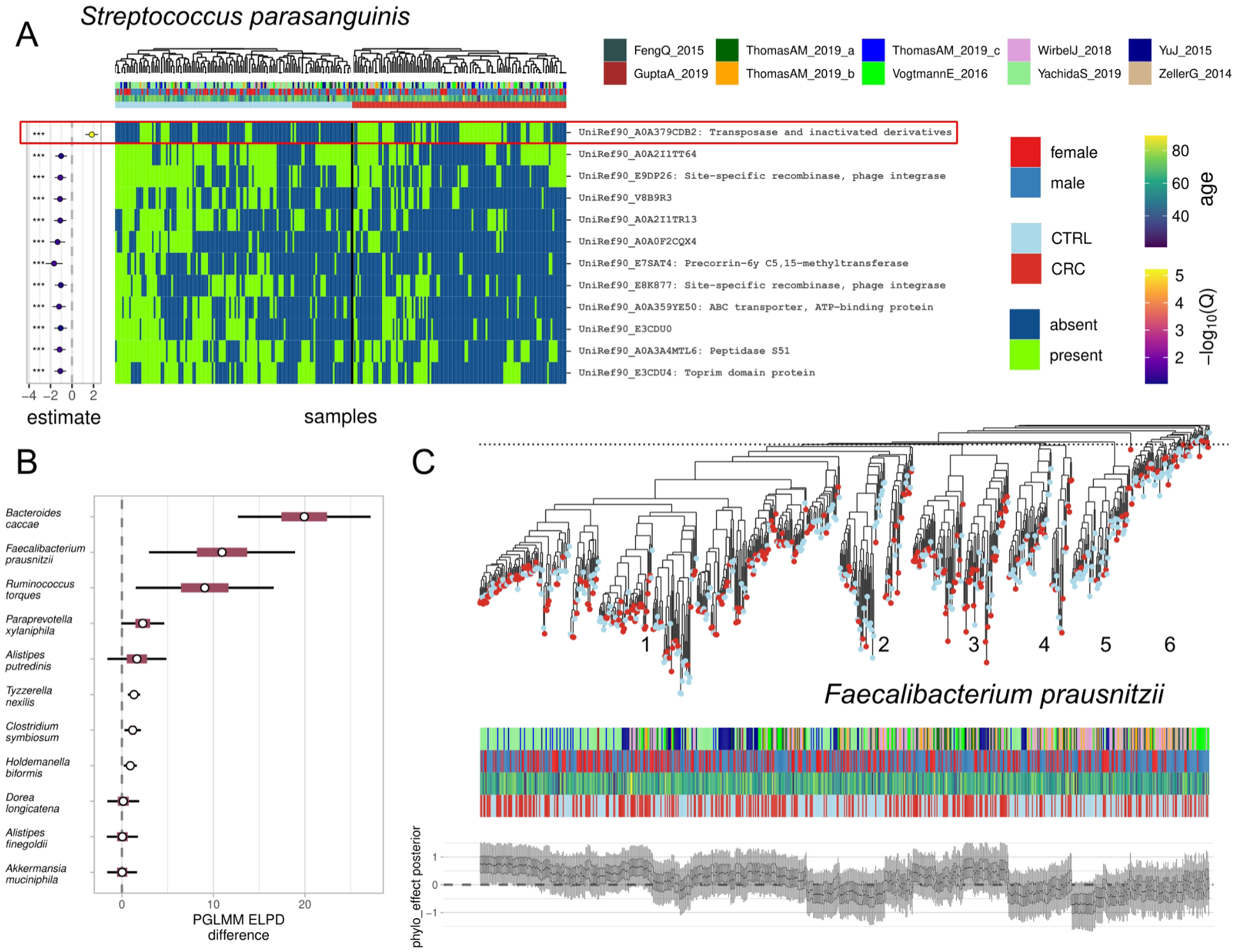
Strain-specific gene carriage and subspecies lineages associated with CRC. **A)** The top 15 strongest gene associations in *Streptococcus parasanguinis* are generally depleted in CRC cases, except for the highlighted gene UniRef90_A0A379CDB2, which is the strongest result in the experiment across all species. Mobile elements are frequently strain-specific, and in the CRC results we indeed observe that many of the top gene hits are related to mobile elements.. **B)** Species with the greatest ELPD difference with respect to CRC, i.e. greatest lineage-specific phenotype enrichments. Phylogeny clearly improves prediction of outcome in all five of the species with the most gene model associations. Inner and outer intervals correspond to 50% and 95% about the normal approximation of the ELPD. **C)** A phylogeny estimated from log copy number profiles (derived from gene profiles) in *Faecalibacterium prausnitzii* shows a strong phylogenetic association with CRC, as evidenced by strong, visible clade effects on CRC risk in the leaf-wise posterior (species wide ELPD difference against base model = 10.9 +/− 4.1). The fine dashed line distinguishes five visually apparent clades and a sixth group of outlying samples. Clades 1 and 3 tend to have higher estimated phylogenetic effects, while groups 2, 4, 5 and 6 have average to low estimates. The model result quantifies the visually apparent pattern of variation in CRC risk by clade and aligns with past reports on the variability of functional potential and phenotypic impact across strains of this species [50].

Returning to the thirteen species with indications of phylogenetic signal, we constructed phylogenies for each by reducing the dimensionality of the gene presence/absence matrices with principal components analysis, then computed the Euclidean distance between samples in this reduced space. The distance matrix was in turn used to approximate phylogeny using neighbor-joining[48]. The resulting trees were used as input to Anpan’s PGLMM model. Several species contained significantly CRC-associated lineages with large leave-one-out predictive performance improvements (**Fig. 6B**). Note that such associations can occur even in species that are not themselves differentially abundant with respect to phenotype; for example, a species may be equally abundant in cases and controls, but consist entirely of one lineage in the former and a second in the latter.

*F. prausnitzii* in particular showed a large, species-wide pattern of different clades exhibiting consistently higher or lower CRC risk (**Fig. 6C**) in a model that also included sex, age, and study as covariates. Among five visually distinguishable sub-clades and a sixth outlying group, clades 1 and 3 typically showed high posterior estimates of phylogenetic effect on CRC risk, while groups 2 and 4-6 exhibited average to low estimates. These visually apparent patterns can be quantified with leave-one-out cross-validated model comparison to a base model with the same covariates but lacking the phylogenetic term. The model comparison reports a clear, strong difference in predictive performance (ELPD improvement of = 10.9 +/− 4.1). This comports with prior reports on both subclades within this species[49] and variability of functional potential and phenotypic impact among those clades[50].

**Figure S3:**
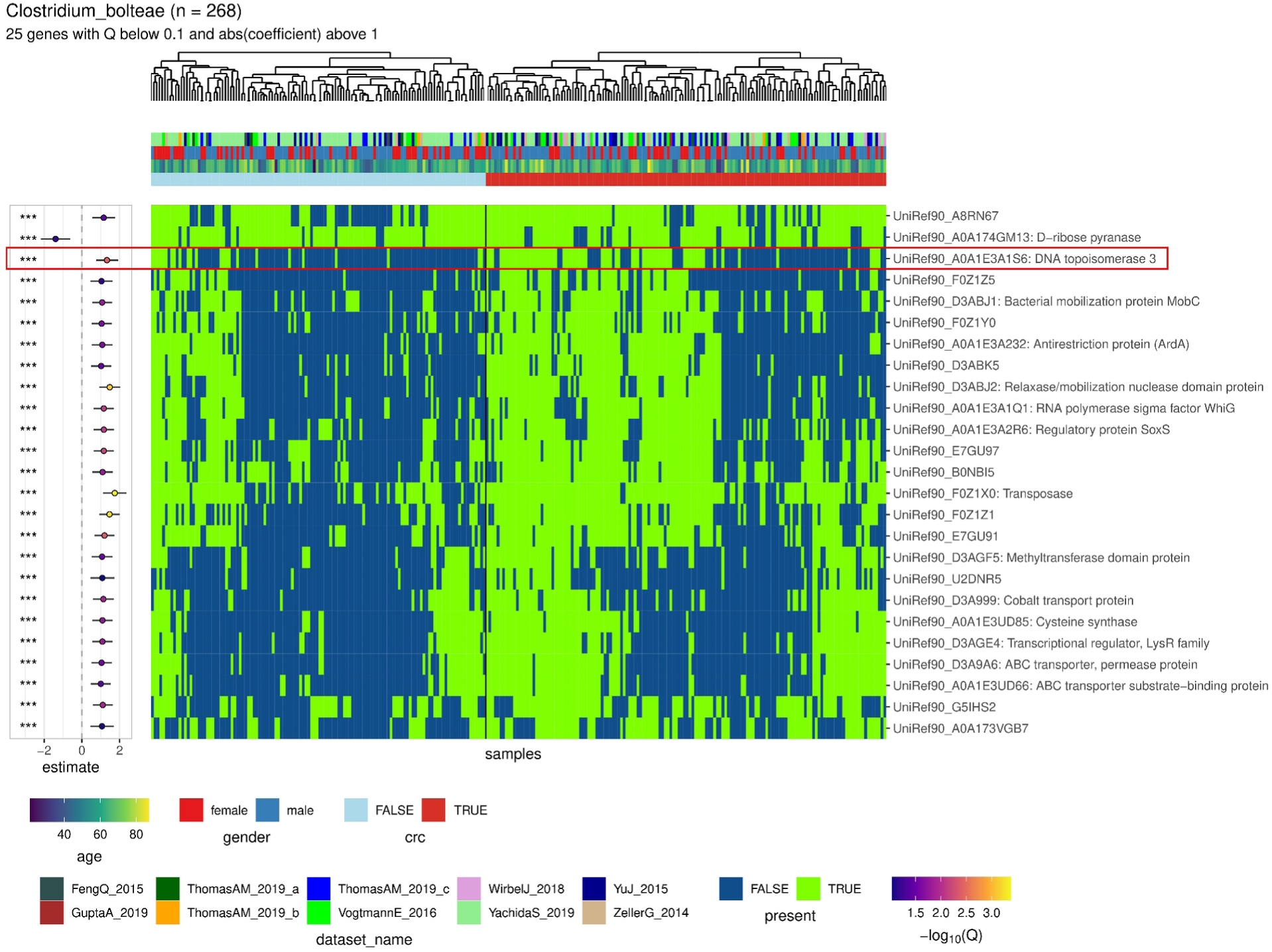
Heatmap results plot of genes in *C. bolteae*. The heatmap shows 25 genes in *C. bolteae* falling withing thresholds Q < .01 and with absolute coefficients greater than 1. Highlighted is UniRef90_A0A1E3A1S6, a topoisomerase more prevalent in CRC cases. This result aligns with the hypothesis that genes enabling genomic restructuring become more prevalent in CRC-derived samples by encouraging adaptation to the inflammatory environment present in the CRC gut.

## Discussion

While there have been many recent advances in strain profiling methods, downstream inferential methods for quantifying strain-level associations with environments, phenotypes, and covariates are limited. To address this, we developed Anpan, a collection of statistical methods for assessing the relationship between microbial strains and outcome variables such as host disease status. These identify three ways in which strains, broadly defined, can be phenotype-associated: gene carriage, pathway retention, and phylogenetic lineage. The gene model uses adaptive filtering to identify which species have sufficient coverage of their genes before using generalized linear models to assess strains’ outcome associations. The phylogenetic model utilizes the correlation structure implied by the relatedness of strains between samples to quantify the contribution of phylogenetic relatedness to phenotype. Finally, the pathway model assesses differences in within-species pathway carriage between experimental groups while accounting for species abundance. Anpan is available as an open-source R package (https://huttenhower.sph.harvard.edu/anpan) alongside extensive documentation and a step-by-step analysis tutorial.

Given the highly multivariate nature of the CRC dataset analyzed here, the associations identified between subspecies structure and CRC status are generally broad patterns that only provide weak indications of the underlying biological mechanism. Non-specific patterns arise from large cohorts such as ours as an unavoidable consequence from the combination weak signals, common processing pipelines, and disparate data collection protocols. Future work in this area will involve focused inspection of the taxa and genetic features identified. Particularly, confirmation of the “mobility for adaptation to inflammation” hypothesis in *S. parasanguinis* and others, as well as characterizing the functional variation in *F. prausnitzii* phylogroups seem like a promising area of research for understanding the interplay of microbial genetics and host health in CRC.

Anpan permits new types of biological hypothesis tests to be applied to microbial communities, as evidenced by our initial investigation of CRC. Microbes which are not among those regularly detected to be differentially abundant during CRC[43, 51] may still preferentially colonize with different phylogenetic lineages during disease (e.g. *F. prausnitzii*) or with distinct gene carriage (e.g. *S. parasanguinis*, *C. bolteae*). Interpretation of these associations requires nuance, particularly since both coding (for the latter) and noncoding (for the former) variants among microbial strains very frequently lack characterization. Even within streptococci, many CRC-specific genes are uncharacterized, as are most of the functional differences among *F. prausnitzii* lineages[50]. Associating these detailed strain-specific features with disease trajectories such as stage or metastasis, however, could very precisely inform the origin of tumor-specific lineage colonization, or the mutual selective pressures exerted by host and microbes in the tumor microenvironment.

Several opportunities to extend the statistical methodology are also plausible. A gene model that accounts for phenomena like linkage and interactions would help increase power and interpretability, as the current model treats genes as independent or as a sparse mixture. This would also bring our analysis of microbial genetics more in line with methods for human genetic epidemiology, although with an even greater need for power and increased sample sizes[52]. Separately, while the existing phylogenetic model allows for fixed effect covariates and offset variables, it cannot incorporate secondary covariance patterns from known sources (e.g. genetic relatedness between geographically proximal cohorts). Realistic datasets commonly feature this sort of secondary covariance, which can make it easy to detect spurious associations from non-biological sources. This could be better-captured by adding a second user-specified covariance term to the model or by algorithmically discounting the components of the phylogenetic covariance matrix that align with the secondary covariates in question.

There are a number of rare but biologically realistic failure modes for the methods presented in this work. When applying the one-at-a-time gene model to a species with distinct subclades that are in turn confounded with the outcome variable, all of the genes unique to the confounded clade will show up as strong hits, regardless of whether they truly affect the outcome. This is akin to the problem of differentiating “driver” versus “passenger” variants in human genetics[53]. A PGLMM fit to a tree derived from the gene profiles can provide evidence that this is occurring, as can diagnostic visualizations of external isolate genome profiles (already implemented in Anpan). The sparsity inherent to the horseshoe gene model can help select the subset of genes most strongly correlated with the outcome, but without external information on the genetic linkage of genes, a sub-clade confounded with the outcome inherently violates the assumptions of the one-at-a-time gene model. It is worth noting that the increasing usage of species definitions based on species genome bins (SGBs), which avoid excessively diverged subclades by design, will help alleviate this issue in the near future. This will be important for poorly characterized species (e.g. rare taxa from environmental samples), for which samples with sufficient coverage of an SGB allow it to be directly placed in the species tree and analyzed as part of a PGLMM.

Even narrowly-defined microbial species can still contain enormous genetic diversity. As a result of the rapid throughput of metagenomic sequencing, new subspecies-level structure is now discovered much faster than individual strains can be experimentally characterized. Downstream inferential methods aimed at quantifying the effects of this variation must therefore utilize within-species aggregation to some degree. The three models of Anpan presented in this work perform the necessary aggregation on interpretable biological units: genes, phylogenies, and pathways. By combining advanced statistical methods, Anpan allows users to quantify the uncertainty in associations between microbial subspecies structure and outcome variables of interest, from cancer risk to environmental properties. This provides much-needed next steps on the path to making microbiome genetic epidemiology both as scalable and as precise as the results that have come to be achieved from human genetics.

## Methods

Anpan provides three core models: a gene model, a phylogenetic model, and a pathway model. The gene model identifies microbial genes that associate with outcome variables such as disease state, and the phylogenetic model identifies within-species phylogenetic patterns of association with outcome variables. The pathway model approaches the inferential framework from the opposite direction, seeking to identify differential pathway abundance between experimental groups.

### Gene model

The gene model uses microbial functional profiles and sample metadata to identify which genes in which species (or other basal clades, e.g. SGBs) best explain sample outcome variables, while accounting for additional covariates like age or gender. There are two key problems when modeling data like this: not all taxa are present in all samples, and when there are enough samples with a given taxon’s genes detectable in them, finding the genes that associate with outcome is very high-dimensional and difficult to adequately power. The gene model in Anpan handles these issues by first applying adaptive sample filtering to classify samples as “well covered” or “poorly covered” for each taxon, so that the modeling can be run only using samples where a species is sufficiently detectable and its gene-level observations accurately detect its genes. The adaptive filtering uses a k-means clustering on two sample statistics - number of nonzero gene observations and median log abundance of nonzero gene observations - to provide this classification.

From there, the gene model applies generalized linear regression models (GLMs, either linear or logistic) that use either FDR correction or a horseshoe prior to classify genes that associate with an outcome beyond what is expected from random variation. As GLMs, they may optionally also include user-supplied covariates to account for additional experimental variables.

#### Gene profile simulations

In order to generate realistic test data for model evaluation, we implemented a simulation framework incorporating components that realistically reflect microbial genetic variation. The simulation framework follows the generative structure of many microbiome studies, where samples are allocated variable abundances of microbial species (which we simulate as log normally distributed with parameters from the SparseDOSSA 2[24] stool preset). Gene log abundances are simulated as a skew-normal(0,1,-1) error added to a linear function of species abundance. Within a species, a small number of genes influence outcome status (simulated as a binary “case” or “control”). Random subjects were generated up to 200 per group (400 in total). Gene log abundance observations were randomly set to zero with probability according to a logistic function centered on the median with growth rate equal to negative one.

### Phylogenetic generalized linear mixed models

Phylogenetic generalized linear mixed models (PGLMMs) are a family of probabilistic models that account for phylogenetic structure. For a PGLMM with a linear regression as the base model, the structure is:

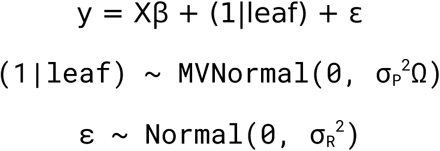

The outcome y is modeled partly as a function of familiar terms from linear regression: covariates X, coefficients ꞵ, and residual noise ε. The key addition is the phylogenetic term (1|leaf), which contains a “random effect” for each observation in y. Unlike typical random effects (which are usually partially pooled toward a global mean but otherwise independent between different levels of the random effect variable), the values in the phylogenetic term follow a pre-specified correlation structure Ω, which is derived from a tree.

The variability of the phylogenetic term is scaled by the “phylogenetic noise” parameter σ_P_. The phylogenetic noise is the key parameter of interest in a PGLMM because it scales the contribution of the tree to the model fit. If the estimate of σ_P_ is close to zero, then the phylogeny does not strongly impact the outcome. In practice, σ_P_ is often contextualized relative to the residual noise σ_R_. There are weak priors placed on σ_P_ and σ_R_ to ensure that the model samples efficiently.

While other methods for accurately fitting PGLMMs exist[54, 55], our implementation provides a number of benefits. First, the Stan model code increases transparency for users, explicitly defining the full set of probabilistic relationships between data and parameters. Second, Stan allows fast, accurate estimation of the full joint posterior distribution of all model parameters. This avoids the pitfalls of inference based on point estimates or assumptions required for approximate algorithms[56], including inaccurate error estimates based on unrealistic assumptions about the curvature of the posterior. Other MCMC algorithms that don’t utilize the gradient of the posterior may fail to converge (and possibly fail to signal to the user the lack of convergence) given the high-dimensionality of the model.

In our experience, our implementation converges quickly (i.e. yields an Rhat convergence diagnostic[57] close to 1) at the default sampler settings and provides highly visible non-convergence warning messages in the rare situation where the model fits badly. The sampler uses the default settings from cmdstanr[58], including four independent chains each with 1,000 warmup samples and 1,000 sampling draws. Additionally, our implementation provides estimates of leaf-wise phylogenetic effects (each individual boxplot in **Fig. 4**), which are usually marginalized out for computational convenience in most other PGLMM implementations. These aid in downstream visualization of results by allowing the juxtaposition of the tree against the posterior on leaf effects (**Fig. 4**) or posterior predictive distributions. This provides users with an intuitive visualization of where in the tree the model is detecting phylogenetic signal, which is not possible with methods that marginalize out leaf-wise effects. Finally, the Stan ecosystem enables the convenient leave-one-out cross-validation that we use for model comparison.

#### Phylogenetic simulations

Simulations for evaluation of Anpan’s PGLMMs covered a range of sample sizes and variance parameters. A single simulation iteration proceeds as follows: We generate a random phylogeny and corresponding correlation and distance matrices using the rtree(), vcv.phylo(), and cophenetic() functions from the ape R package[48], respectively. A phylogenetically determined outcome variable is generated by adding a random multivariate normal draw scaled by σ_P_ to a standard normal draw scaled by σ_R_.

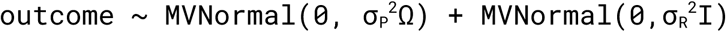

We then fit two variations of PERMANOVA using the adonis2() function from the vegan package and MiRKAT() from the MiRKAT package[59]. We also fit unregularized and regularized (the latter adds a Gamma(1,2) prior on the ratio of σ_P_ to σ_R_) PGLMMs using Anpan. We vary the number of tips in the phylogeny, σ_P_, and σ_R_ across a grid of six unique values per parameter from 10 to 450, 0 to 2.09, and 0 to 2.09, respectively. We run 30 simulation iterations per grid point at each of the 210 grid points (grid points with σ_P_ and σ_R_ both equal to zero are dropped).

### Pathway model

Anpan’s pathway model is constructed using random effects to capture differential pathway carriage (i.e. slope) per pathway per taxon. For a consistently-carried pathway, pathway presence and taxon abundance will be perfectly correlated, although their abundance distributions may differ between phenotypes. Anpan looks for pathways that differ substantially in abundance between phenotype groups while accounting for the correlation with species abundance. Using standard random effects model syntax[60], the resulting formula is:

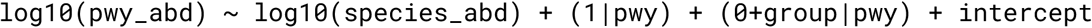

There are a few important features to note about this model. First, because the model works with log abundances, zeros are discarded. During development and evaluation, models that included a censoring feature for zero observations as unknown low values added significant computational complexity while making no practical difference to results. Second, the relationship between species abundance and pathway abundance is measured globally using data from all pathways in the taxon. Third, each pathway has its own intercept (the (1|pwy) term), but these intercepts are partially pooled across pathways. The prior on these intercepts is a Normal(0, σ_int_) distribution. The prior on σ_int_ is a t(5, 0, 2.5) distribution. Fourth, each pathway has its own group effect (the (0+group|pwy) term). These are also pooled across pathways, but they are independent of the pathway intercepts (signaled by the 0 in the formula). These terms come from a Normal(0, σ_effects_) distribution. The prior on σ_effects_ is a relatively aggressive exponential(3) distribution, which encodes the regularizing prior belief that very few pathways will differ substantially between conditions.

Finally, there are some additional weak priors on the global intercept, fixed effects, and error term. Specifically, these are, respectively, a t(5, EM, 2.5) distribution (where EM is the empirical mean of the log10 pathway abundances), a Normal(1,1) distribution, and a t(3,0,2.5) distribution. These ensure that the model is identifiable and samples efficiently without strongly influencing the result.

The model is fit with Stan and typically samples efficiently, yielding high effective sample sizes and Rhat values[57] close to one. Pathways are called as “hits” by three (configurable) criteria: 1) the 98% posterior intervals on the pathway:group effect excludes zero, 2) the absolute posterior mean exceeds 0.25 (i.e. a 1.77-fold increase or decrease), and 3) the estimated relationship between species_abundance and pathway abundance is positive.

### Colorectal cancer data and analysis

CRC data were derived from eight publicly available metagenomic studies: Yu et al.[37] (n = 53 controls, n = 75 cases), Feng et al.[38] (n = 61 controls, n = 46 cases), Zeller et al.[39] (n = 61 controls, n = 53 cases), Vogtmann et al.[40] (n = 52 controls, n = 52 cases), Gupta et al.[41] (n = 30 controls, n = 30 cases), Thomas et al.[42] (n = 89 controls, n = 97 cases), Wirbel et al.[43] (n = 65 controls, n = 60 cases), and Yachida et al.[44] (n = 251 controls, n = 187 cases). Observations from some studies were dropped due to intermediate phenotypes (e.g. adenoma, CRC stage zero) or incongruous phenotypes (metastases in control patients).

Data were acquired using the curatedMetagenomicData R package[61]. The data were used as pre-processed by this pipeline, which includes manual metadata curation and profiling using MetaPhlAn and HUMAnN[36]. Starting from the HUMAnN gene profiles, each taxon-specific gene profile is filtered in three steps: an initial prevalence filter, adaptive k-means filtering, and a final prevalence filter. The two prevalence filters remove genes that do not occur fewer than floor(0.005*N) (six in the case of our eight combined CRC datasets) times across all subjects or more than N - floor(0.005*N) times(1,256 in this case). The adaptive k-means filter fits a k-means classifier with k=2 to the sample-wise data of two sample statistics: number of non-zero observations and median raw abundance in the gene profile. The adaptive filter then discards samples in the cluster with few non-zero observations and low abundance. After the three filtering steps, gene profiles are binarized into zeros and non-zeros to act as indicators of gene presence. These filtered, binarized profiles are the input to the modeling step.

To construct the phylogenetic trees used in this work, we estimated principal components of the filtered gene presence tables. This was done in order to reduce the impact of random noise in the gene profiles on tree estimation. Due to the inconsistent complexity of genetic structure between different taxa, rather than take a fixed number of principal components, we used an adaptive number of principal components. We compared the relative magnitude of the eigenvalues of the decomposition of the filtered gene profile and selected up to the first eigenvalue that was under one tenth of the magnitude of the overall first eigenvalue. After projecting the gene profile onto this number of principal component dimensions, we computed Euclidean distances between samples then used neighbor-joining to compute a tree (using the nj() function from the ape R package[62]). Finally, the tree topology was reorganized using the ladderize() function from ape.

## Declarations

### Code availability

The open source R and Stan package is available from https://huttenhower.sph.harvard.edu/anpan, along with installation instructions and a detailed walkthrough vignette.

### Data availability

CRC data are previously published[37–44]. Metagenomes were retrieved from the ENA and processed uniformly using the bioBakery 3 metagenome workflow[36].

### Funding

This work was funded by NIH NIDDK R24DK110499 (CH), the Cancer Grand Challenges Team OPTIMISTICC (CH), and a Strategic Research Alliance between Astellas Pharmaceuticals Inc. and Harvard University(CH).

### Author Contributions

Andrew R. Ghazi: Conceptualization, Methodology, Software, Validation, Investigation, Writing - Original Draft, Writing - Review & Editing, Visualization

Yan Yan: Data Curation, Writing - Review & Editing

Kelsey N. Thompson: Data Curation, Validation, Writing - Review & Editing

Zhendong Mei: Validation, Writing - Review & Editing

Amrisha Bhosle: Validation, Writing - Review & Editing

Fenglei Wang: Validation, Writing - Review & Editing

Kai Wang: Validation, Writing - Review & Editing

Eric A. Franzosa: Conceptualization, Methodology, Writing - Review & Editing, Supervision

Curtis Huttenhower: Conceptualization, Writing - Review & Editing, Funding acquisition

### Competing Interests

The authors have no competing interests to declare.

## Supporting information

Supplemental Figuren 3

## Notes

https://github.com/biobakery/anpan

### Competing Interest Statement

The authors have declared no competing interest.

