## Supplementary figures and images for "Quantifying Metagenomic Strain Associations from Microbiomes with Anpan"

### Supplemental Figure 1

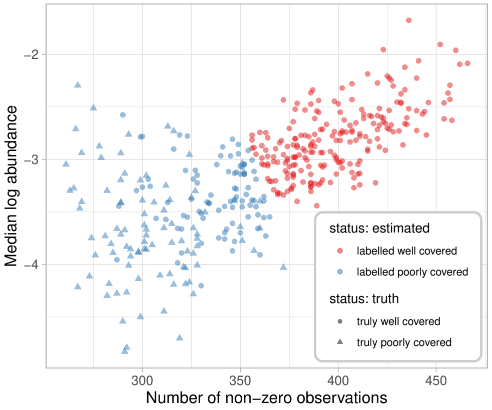

### Supplemental Figure 2

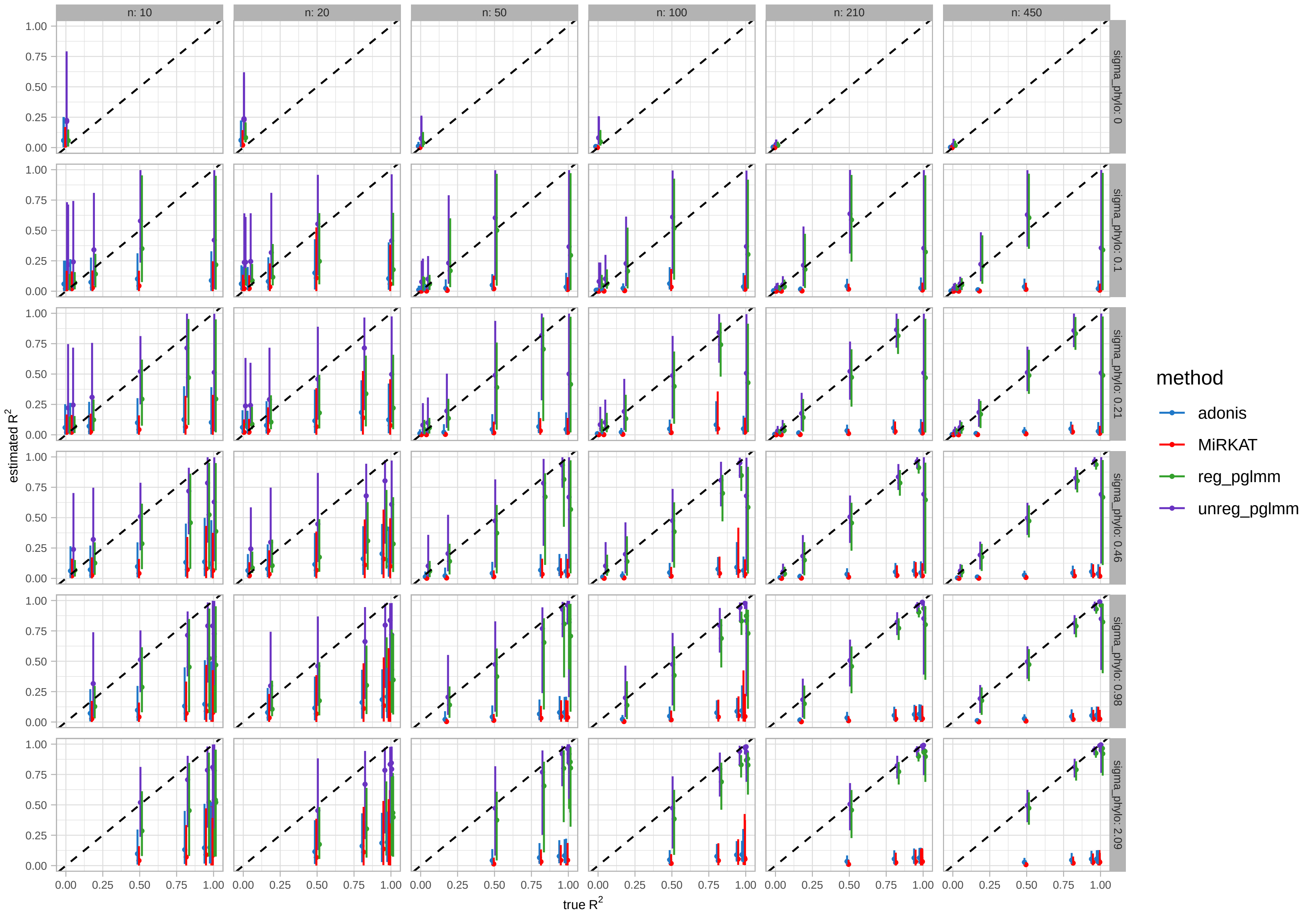

### Supplemental Figuren 3

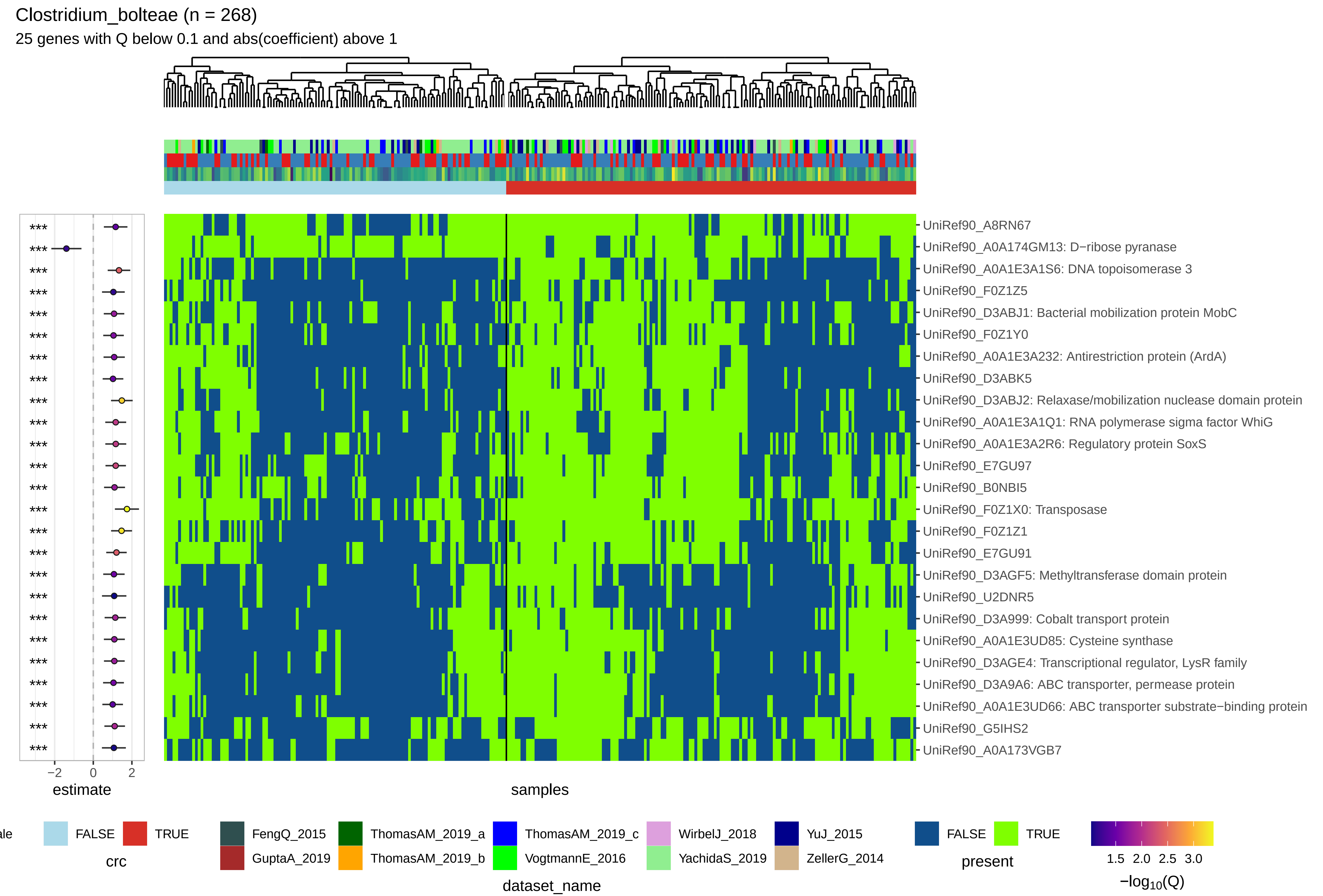
